# Graphtyper: Population-scale genotyping using pangenome graphs

**DOI:** 10.1101/148403

**Authors:** Hannes P. Eggertsson, Hakon Jonsson, Snaedis Kristmundsdottir, Eirikur Hjartarson, Birte Kehr, Gisli Masson, Florian Zink, Aslaug Jonasdottir, Adalbjorg Jonasdottir, Ingileif Jonsdottir, Daniel F. Gudbjartsson, Pall Melsted, Kari Stefansson, Bjarni V. Halldorsson

## Abstract

A fundamental requisite for genetic studies is an accurate determination of sequence variation. While human genome sequence diversity is increasingly well characterized, there is a need for efficient ways to utilize this knowledge in sequence analysis. Here we present Graphtyper, a publicly available novel algorithm and software for discovering and genotyping sequence variants. Graphtyper realigns short-read sequence data to a pangenome, a variation-aware graph structure that encodes sequence variation within a population by representing possible haplotypes as graph paths. Our results show that Graphtyper is fast, highly scalable, and provides sensitive and accurate genotype calls. Graphtyper genotyped 89.4 million sequence variants in whole-genomes of 28,075 Icelanders using less than 100,000 CPU days, including detailed genotyping of six human leukocyte antigen (HLA) genes. We show that Graphtyper is a valuable tool in characterizing sequence variation in population-scale sequencing studies.

## Introduction

Advances in DNA sequencing technology have improved characterization of sequence diversity in the human genome and have resulted in refinements of the reference sequence^1–4^. The human reference sequence is extremely useful, but it represents a consensus of genomes and therefore it does not capture sequence variation within or between populations^5,6^.

In the latest version of the human reference genome (GRCh38), there are several alternate loci where the sequence variation is too complex to be represented with a single sequence. These loci are generally highly polymorphic, and many are known to co-segregate with disease and are therefore of great interest in population genetics. The most prominent example, the human leukocyte antigen (HLA) region, is known to associate with a number of immune mediated human diseases^7^. Given the importance of this region, it has been further characterized in the IPD-IMGT/HLA database^8^, which contains a large collection of known HLA allele sequences. Such variation should be included in genome diversity analyzes.

Short-read sequencing is the standard in genome-wide sequence analysis. Most common approaches for discovering sequence variants involve aligning sequence reads to a reference genome^9^ and searching for variants as alternative sequences in read alignments (Figure 1a i). However, some reads cannot be aligned to a reference genome, particularly those originating from highly polymorphic regions and regions absent from the reference genome. Reference genome alignments are also generally done without awareness of variation, causing mapping bias towards the reference allele and misalignments around indels^10,11^.

**Figure 1:**
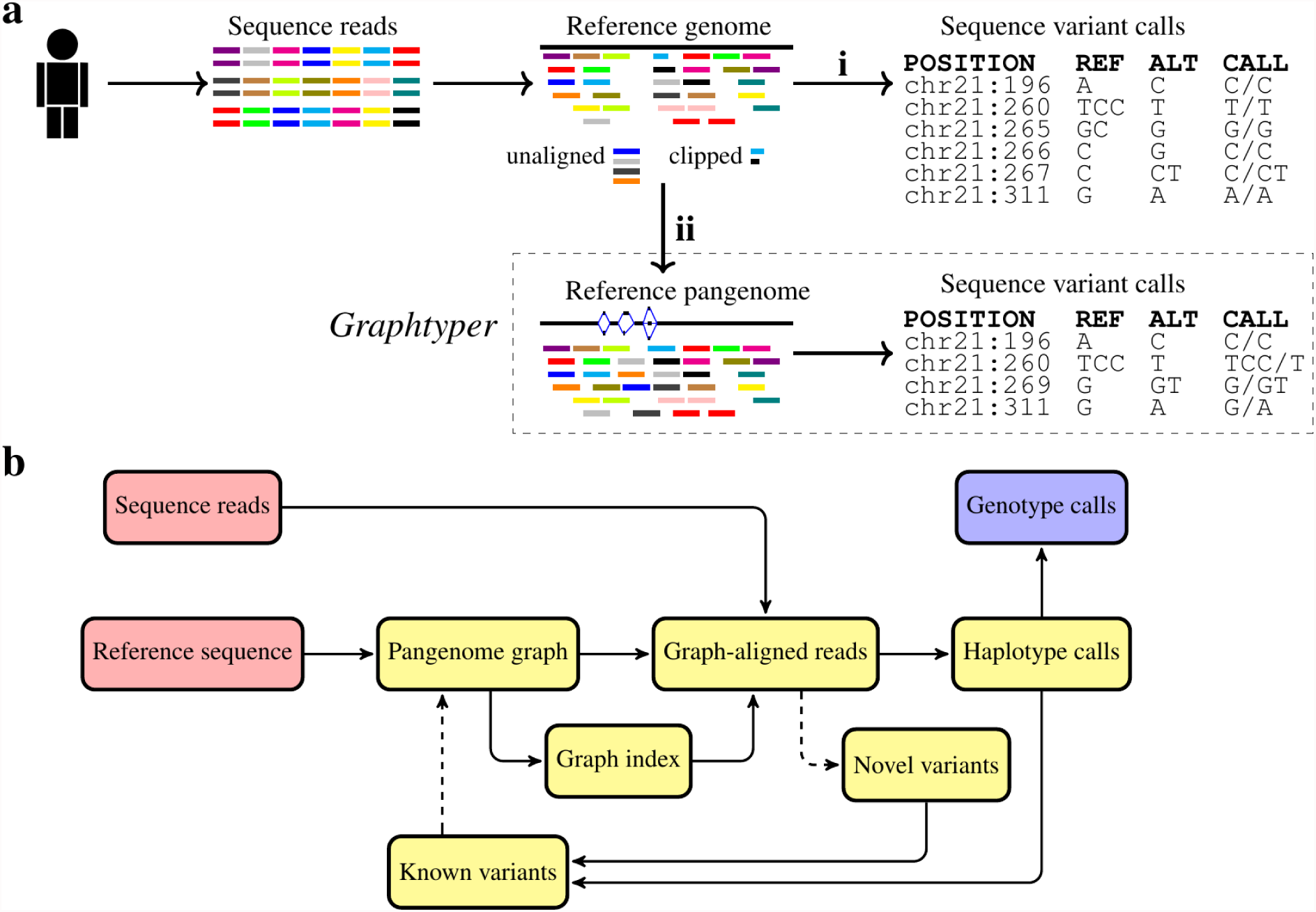
**(a)** Overview of two genotyping pipeline designs. **(i)** A commonly used genotyping pipeline, where sequence reads are aligned to a reference genome sequence and sequence variant calls are made from sequence discordances between the sequence reads and the reference sequence. **(ii)** Graphtyper’s genotyping pipeline. Sequence reads are realigned to a variant-aware pangenome graph and variants are called based on which path the reads align to. **(b)** Graphtyper’s iterative genotyping process. Dashed paths are optional. As input, Graphtyper requires a reference genome sequence and sequence reads (red) and outputs genotype calls (blue) of variants.

Richer data structures that utilize the large amount of available sequence variation data promise to alleviate some of the limitations of previous methods^12–15^. Although approaches that find polymorphisms in reference-free assemblies have been developed to avoid these limitations^16,17^, *de novo* assembly algorithms remain computationally expensive, have less sensitivity^17^, and use data structures that have a complex coordinate system.

Pangenomes^12,18,19^ have recently been proposed to counter weaknesses of both reference alignments and *de novo* assemblies by extending the linear reference alignments with variation-aware alignments^20^. Pangenomes incorporate prior information about variation, allowing read aligners to better distinguish between sequencing errors in reads and true sequence variation. Unlike *de novo* assembly algorithms, pangenomes represent sequence variation with respect to the reference genome, enabling a direct access to its annotated biological features. Variation-aware data structures, such as pangenomes, also allow read mapping and genotype calling to be performed in a single step^12^.

Graph-like data structures with directed edges have commonly been used to represent pangenomes^19,21–24^. In an idealized pangenome graph, nodes represent sequences and the sequence of every genotyped individual genome is a path in the graph, but not necessarily vice versa. A number of algorithms have recently been developed that tackle the problems of graph construction, indexing and alignment of sequence reads to graphs ^19,21,25–27^, Paten *et al*.^24^ provide a recent survey of current efforts. However, there is no method that combines these operations and uses the resulting alignments to update the graph with novel variation for the purpose of variant calling^12^.

Here we present Graphtyper, a method and software for discovering and genotyping sequence variants in large populations using pangenome graphs. Graphtyper realigns all sequence reads of a genomic region, including unaligned and clipped sequences, to a variation-aware graph (Figure 1a ii). Concomitantly, it aligns sequence reads and genotypes sequence variants present in its graph. Furthermore, Graphtyper discovers novel single nucleotide polymorphisms (SNPs) and short sequence insertion or deletion variants (indels), which can be used to update the pangenome graph (Methods).

An important benefit of Graphtyper’s realignment step is to improve read alignments near indels. Figure 2a shows how Graphtyper represents three common sequence variants, a 40-bp deletion and two SNPs. Using variation-aware realignment, Graphtyper is capable of better characterization of the region’s variation than previous methods, with no Mendelian errors (Figure 2b) and no falsely reported additional sequence variants around the indel (Supplementary Table 1) due to misaligned sequence reads (Supplementary Figure 1).

**Figure 2:**
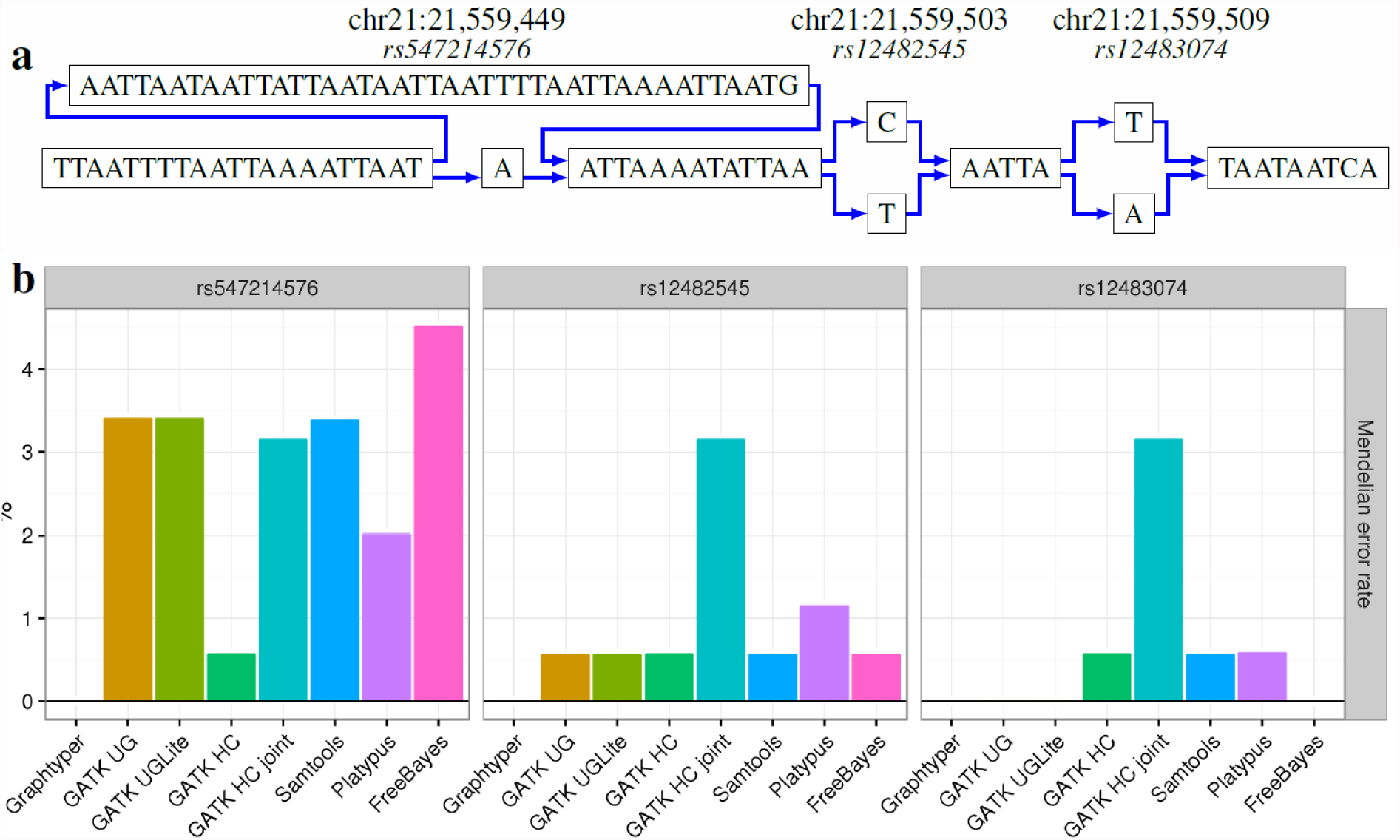
**(a)** The genomic region chr21:21,559,430-21,559,518 (GRCh38) and three previously reported sequence variants represented with a pangenome graph. **(b)** Mendelian error rates of the three previously reported sequence variants called by eight genotypers. The Mendelian error rate is measured in 230 Icelandic parent-offspring trios.

## Results

### Data structures and genotyping pipeline

Graphtyper uses a reference sequence and optionally all known sequence variants as input to construct pangenome graphs. Sequence reads mapped to a genomic region of the reference sequence, including unaligned and trimmed reads, are realigned to the pangenome graph. Using these graph alignments, Graphtyper discovers variants within the genomic region. This process is iterated several times (Supplementary Note 4), i.e., a pangenome graph is constructed, indexed and aligned with sequence reads, from which novel variants are discovered and previously discovered variants are genotyped (Figure 1b).

The underlying pangenome data structure is a directed acyclic graph (DAG) where edges connect nodes that contain a DNA sequence (Supplementary Note 1). Graphtyper takes as input a reference genome and a list of known variants. Each known variant is a record of a chromosomal position, a reference allele, and one or more alternative alleles. First, variant records with overlapping reference alleles are merged into a single record (Figure 3a). Second, *allele nodes* are constructed, containing the sequence and start position of each allele of the variant records. Third, *reference nodes* are constructed between two adjacent variant records, storing the corresponding reference sequence and its start position. Finally, nodes at adjacent positions are connected. Paths in the graph alternate between *reference* and *allele* nodes and nodes that share a start position are parallel to each other. Each character in an allele node sequence is given a position equal to the first position of the node plus the character’s offset from that position (Figure 3b). Allele node positions longer than the reference allele are assigned new unique positions (*z*_1_ and *z*_2_ in Figure 3b) to avoid conflicts with the following positions. The final graph represents the reference sequence and all haplotypes in the population as paths.

**Figure 3:**
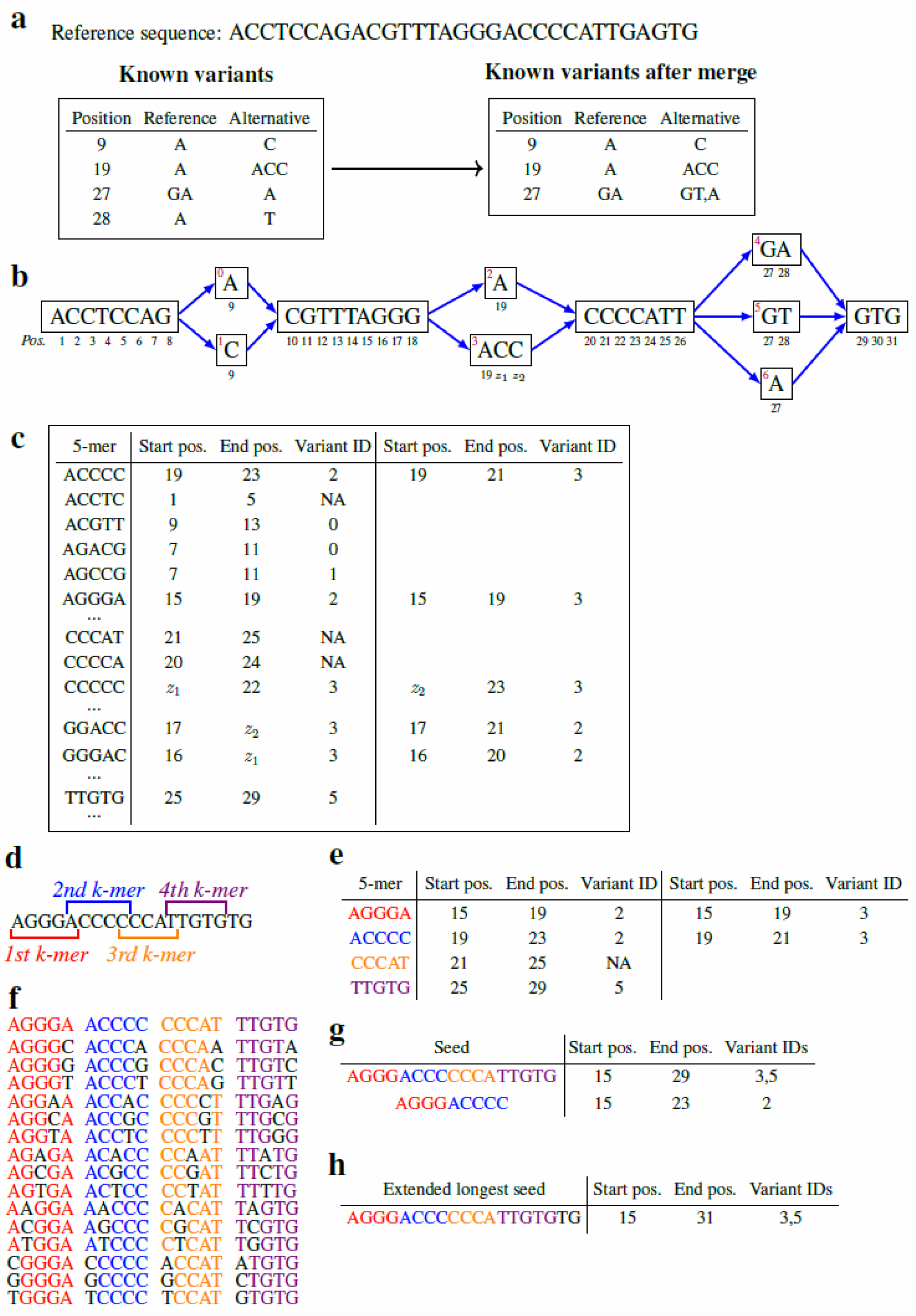
**(a)** An example reference sequence and its known variation. All overlapping variants are merged. **(b)** Constructed pangenome reference graph. We draw the path of the reference sequence as the topmost path. **(c)** The index data structure with k = 5. 5-mers in the graph are mapped to a list of its start position, end position, and a variant ID which it overlaps, if any. **(d)** Four k-mers are extracted from a sequence read. Each k-mer overlaps its neighbor k-mer by one character. **(e)** An example look-up of the k-mers from the index data structure from c). **(f)** All extracted k-mers with a single substitution. **(g)** Seeds are generated from matches in the index look-up. **(h)** Final graph alignment after extending the longest seed.

Aligning sequence reads by traversing the graph is time consuming. To expedite graph alignments, the graph structure is preprocessed by creating an index that maps *k*-mers to their start and end positions in the reference genome and to overlapping allele nodes (if any) (Figure 3c, Methods). Read alignment then follows the seed-and-extend paradigm (Figure 3d-3h, Methods, and Supplementary Note 2).

The output of each iteration is a file in variant-call format (VCF) including both newly and previously discovered variants, which Graphtyper uses to update the graph in the next iteration (Methods).

### Population-scale genotyping

We compared Graphtyper to seven widely used genotyping pipelines on human chromosome 21 in a set of 691 whole-genome sequenced Icelanders (Table 1). Of these, 404 individuals were contained in 230 trios (parent-offspring trio families). The genotypers used were Genome Analysis ToolKit UnifiedGenotyper (GATK UG)^28^, GATK-Lite UnifiedGenotyper (UGLite), GATK HaplotypeCaller (HC), GATK HC GVCF joint genotyping (HC joint), Samtools^29^, Platypus^17^, and FreeBayes^30^ (Supplementary Note 4). Known sequence variants were not given to Graphtyper as input, all pipelines were given the same BAM files and reference sequence (GRCh38).

**Table 1:** Raw sequence variant calls comparison of 691 whole-genome sequenced Icelanders of human chromosome 21.

| Genotyping pipeline | GraphTyper | GATK UG | GATK UGLite | GATK HC | GATK HCjoint | Samtools | Platypus | FreeBayes |
| --- | --- | --- | --- | --- | --- | --- | --- | --- |
| Sequence variant records | 453,288 | 451,131 | 451,415 | 311,731 | 418,949 | 411,907 | 424,000 | 596,499 |
| SNPs | 406,087 | 397,821 | 397,890 | 267,949 | 352,293 | 336,544 | 301,066 | 562,319 |
| Transitions/Transversions | 1.49 | 1.46 | 1.46 | **1.75** | 1.56 | 1.50 | 1.38 | 0.70 |
| Indels | 47,866 | 53,310 | 53,525 | 46,779 | 73,934 | 75,363 | 110,347 | 33,086 |
| MNPs | 1,002 | 0 | 0 | 0 | 0 | 0 | 26,086 | 21,044 |
| Complex | 3,682 | 0 | 0 | 0 | 34,592 | 0 | 0 | 4,532 |
| Common (dbSNP b149) | 157,288 | **158,700** | 158,590 | 153,543 | 158,411 | 157,998 | 156,280 | 136,882 |
| SNPs | 145,143 | 145,723 | **145,724** | 140,533 | 144,858 | 145,135 | 142,417 | 126,653 |
| Indels | 12,145 | 12,977 | 12,866 | 13,010 | 13,553 | 12,863 | **13,863** | 10,229 |
| Alternative alleles called in trios | 454,157 | 447,144 | 450,241 | 312,275 | 435,511 | 392,960 | 408,648 | 448,429 |
| Germline<sub>estimated</sub> | **267,057** | 264,447 | 264,753 | 237,978 | 254,427 | 255,630 | 228,646 | 200,776 |
| FDR<sub>estimated</sub> | 41.20% | 40.86% | 41.20% | 23.79% | 41.58% | 34.95% | 44.05% | 55.23% |
| SNPs | 371,214 | 366,068 | 366,019 | 243,815 | 307,024 | 295,707 | 255,775 | 364,942 |
| Germline<sub>estimated</sub> | **232,256** | 227,858 | 227,872 | 206,084 | 216,448 | 215,042 | 183,375 | 172,226 |
| Non-SNPs | 82,943 | 81,076 | 84,222 | 68,460 | 128,487 | 97,253 | 152,873 | 83,487 |
| Germline<sub>estimated</sub> | 34,801 | 36,589 | 36,881 | 31,894 | 37,979 | 40,588 | **45,271** | 28,550 |
| Common dbSNP calls |  |  |  |  |  |  |  |  |
| Mean transmission rate | 49.98% | 50.08% | 50.08% | 50.01% | 50.01% | 50.11% | 49.47% | 50.17% |
| Mean missing call rate in trios | **0.201%** | 0.290% | 0.289% | 0.329% | 0.252% | 0.375% | 0.445% | 0.259% |
| Mendelian accuracy | **99.52%** | 99.48% | 99.48% | 99.37% | 99.41% | 99.38% | 99.11% | 99.44% |
| Recalled microarray calls | 3,188,286 | 3,197,365 | 3,197,365 | 3,189,368 | 3,193,870 | **3,200,590** | 3,056,002 | 2,640,485 |
| Concordance | 99.79% | 99.80% | 99.80% | 99.78% | 99.76% | 99.78% | 99.20% | **99.90%** |
| Only ref/ref array calls | 99.92% | 99.92% | 99.92% | 99.93% | 99.93% | 99.93% | 99.90% | **99.96%** |
| Only ref/alt array calls | 99.65% | 99.63% | 99.63% | 99.54% | 99.52% | 99.58% | 99.01% | **99.83%** |
| Only alt/alt array calls | 99.71% | 99.81% | 99.81% | 99.80% | 99.74% | 99.76% | 97.94% | **99.85%** |
| CPU time [hr] | 582 | **576** | 1,640 | 12,964 | 1,216 (87<sup>*</sup>) | 594 | 3,173 | 1,030 |
| Time per sample [hr] | 0.842 | **0.834** | 2.373 | 18.761 | 1.76 (0.13<sup>*</sup>) | 0.860 | 4.592 | 1.491 |
| Mean memory [GB] | 10.68 | 50.17 | 40.55 | 65.22 | 51.98 | **1.97** | 6.31 | 6.77 |
| Maximum memory [GB] | 45.40 | 52.72 | 45.86 | 307.47 | 53.58 | **2.69** | 50.15 | 196.03 |

Our results show that GATK UG, Graphtyper and Samtools all had comparable compute times and completed the genotyping in between 576 and 594 hours (Table 1). The other five genotypers required considerably greater compute times (1,030-12,964 hours).

We assessed the raw output of all eight genotyping pipelines to compare them independent of filtering technique and to include analysis of all germline variation, somatic variation, and wrongfully reported variation due to sequencing or alignment errors. Compared to other genotypers, Graphtyper called a large number of SNPs (406,087) with a reasonably high ratio of transitions (Ti) to transversions (Tv) (1.49). We observed that all eight genotypers had a large excess of alternative alleles with a transmission rate below 50% (Supplementary Figure 2). We also observed higher Ti/Tv ratios among alleles with higher transmission rates (Supplementary Figure 3). Motivated by these realizations, we estimated the number of germline alternative alleles based on the transmission rate of the alternative alleles in the 230 trios (Methods). Graphtyper detected the largest number of estimated germline alternative alleles in the trios (267,057), followed by GATK UGLite (264,753) and GATK UG (264,447) (Table 1).

We found 105,302 SNPs and 7,694 indels that were called by all eight genotypers and have been reported as common (minor allele frequency > 1% in any population) in dbSNP build 149. In the 230 trios, Graphtyper called these sequence variants with a mean transmission rate of 49.98%, very close to the expected 50%. Graphtyper had the highest Mendelian accuracy (99.52%) and the lowest number of missing genotype calls (0.201%) (Table 1). We also compared SNP calls to our in-house microarray genotypes (Methods), all genotyping pipelines were highly concordant (>99%).

From our comparison of genotypers, we concluded that Graphtyper and GATK UG were the two best genotypers for population-scale genotyping in terms of performance, accuracy and sensitivity. We assessed a call set of highly confident Graphtyper sequence variants using our own filtering criteria and filtered the GATK call sets (UG, HC and HC joint) using their available ‘best practices’ filtering criteria (Supplementary Note 4). Graphtyper achieved substantially lower estimate of false discovery rate (FDR) (2.19%) than the other call sets (10.26-31.22%), but also had lower estimated number of germline alternative alleles (200,984) than the other call sets (214,801-240,020) (Supplementary Table 2).

We measured their scalability by genotyping chromosome 21 on a dataset of 15,220 Icelanders, in which there are 1,729 trios (3,863 unique individuals). Our results show that Graphtyper scales much better than GATK UG (Figure 4), with GATK UG using approximately 2.5x more time for computations than Graphtyper (Table 2). The compute time used by Graphtyper per sample did not increase substantially when the sample size increased from 691 to 15,220 (changed from 0.842 hr/sample to 0.867 hr/sample), while GATK UG used 2.65x more compute time per sample (changed from 0.834 hr/sample to 2.206 hr/sample).

**Figure 4:**
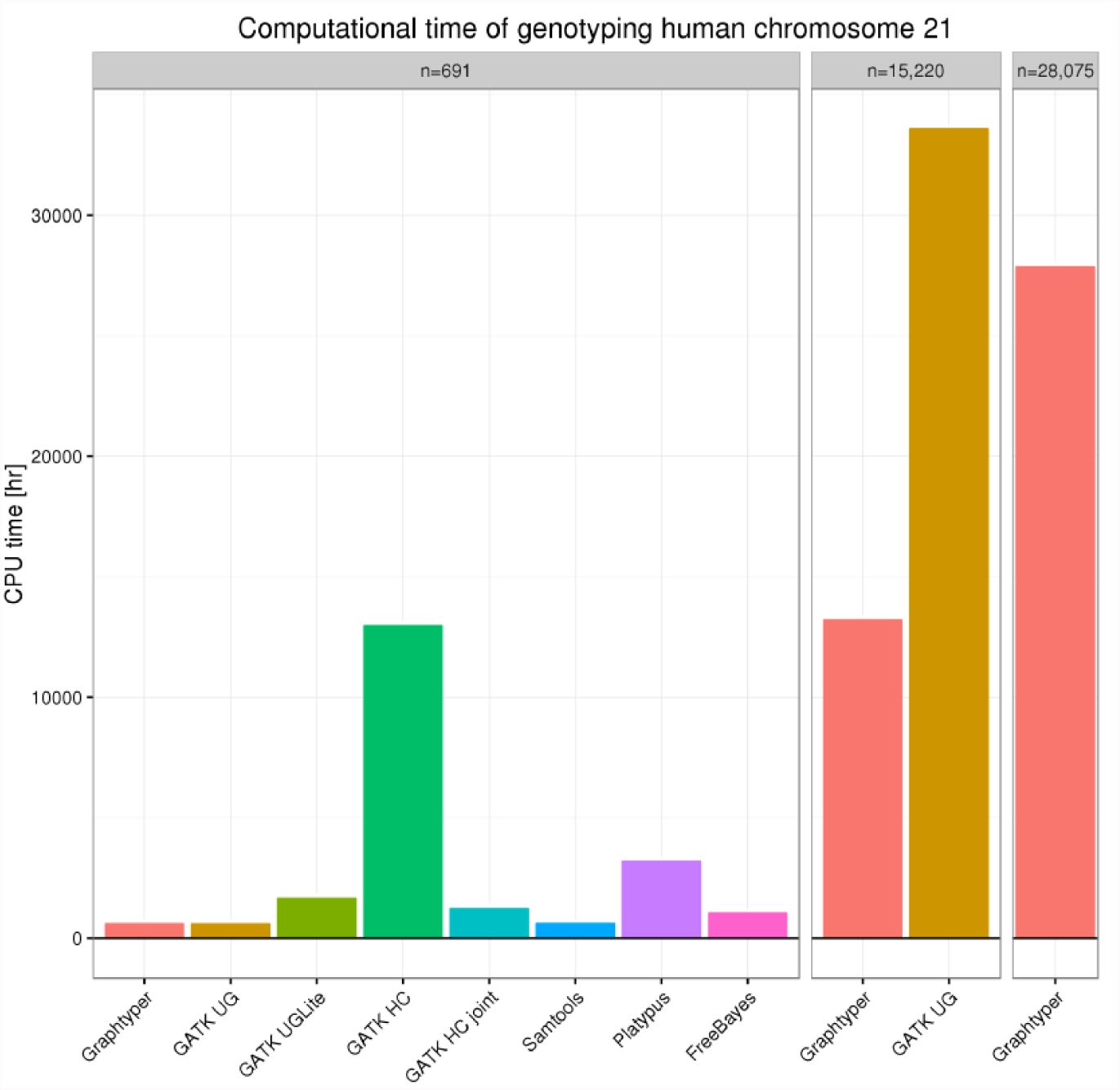
Comparison of compute times required to genotype chromosome 21 on three whole-genome sequence datasets.

**Table 2:** Comparison of Graphtyper and GATK UG genotyping chromosome 21 of 15,220 sequenced Icelanders.

| Genotyping pipeline | Raw |  | Filtered |  |
| --- | --- | --- | --- | --- |
|  | Graphtyper | GATK UG | Graphtyper | GATK UG |
| Sequence variant records | 1,101,540 | 1,160,333 | 473,813 | 493,620 |
| SNPs | 1,024,677 | 1,035,206 | 437,844 | 423,407 |
| Transitions/Transversions | **1.14** | 1.06 | 2.24 | 2.27 |
| Indels | 81,848 | 125,127 | 36,086 | 70,213 |
| MNPs | 3,487 | 0 | 133 | 0 |
| Complex | 10,707 | 0 | 888 | 0 |
| Alternative alleles called in trios | 979,451 | 1,032,839 | 338,266 | 394,679 |
| Germline<sub>estimated</sub> | 383,998 | **397,283** | **308,204** | 305,404 |
| FDR<sub>estimated</sub> | **60.79%** | 61.53% | **8.89%** | 22.62% |
| SNPs | 821,098 | 850,761 | 304,881 | 294,004 |
| Transitions/Transversions | **1.01** | 0.92 | 2.18 | 2.19 |
| Germline<sub>estimated</sub> | 340,313 | **349,878** | **281,972** | 264,441 |
| FDR<sub>estimated</sub> | **58.55%** | 58.87% | **7.51%** | 10.06% |
| Non-SNPs | 158,353 | 182,078 | 33,385 | 100,675 |
| Germline<sub>estimated</sub> | 43,685 | **47,405** | 26,232 | **40,963** |
| FDR<sub>estimated</sub> | **72.41%** | 73.96% | **21.43%** | 59.31% |
| CPU time [hr] | **13,192** | 33,573 | — | — |
| Time per sample [hr] | **0.867** | 2.206 | — | — |

Based on the transmission of alternative alleles the 1,729 trios, we observed that the FDR increased for Graphtyper and GATK UG compared to the 230 trio dataset in both raw and filtered call sets. We estimated that Graphtyper detected more germline alternative alleles (308,204) with a significantly lower FDR (8.89%) than GATK UG (305,404 and 22.62%, respectively) in the filtered call sets (Table 2).

### Single sample genotyping

We assessed the single sample genotyping performance of Graphtyper on a well-studied parent-offspring trio (NA12878, NA12891 and NA12892). Whole-genome sequence data (50x 101-bp paired-end Illumina HiSeq 2000) of these samples are publicly available through the Platinum Genome project^31^. We genotyped each sample independently using the same genotyping pipelines as in our population-scale experiment. We ran Graphtyper with and without initializing its graph structure with publicly available common (minor allele frequency > 1% in any population) sequence variants (dbSNP build 150).

We assessed sequence variant call sets of the offspring (NA12878) by comparing it to the set of publicly available high-confidence variant calls^31^ to measure variant recall rate and precision. Based on the genotyping of the parents (NA12891 and NA12892), we estimated FDR and the number of transmitted germline alternative alleles in the trio (Methods).

Our results show that even without the knowledge of known variation, Graphtyper has a considerably better recall rate (98.14%) than the other genotypers (90.24-95.91%), high precision (99.774%), and overall the highest number of validated calls (4,081,193) (Table 3). As expected, the knowledge of common dbSNP variants increased Graphtyper’s recall rate (to 98.46%), in particular at non-SNP sites where it increased from 91.23% to 93.38%. Consistent with its measured high recall rate, we also estimated that Graphtyper called the highest number of germline alternative alleles in the trio (5,991,012 and 5,874,556 with and without dbSNPs, respectively), substantially more than the other genotypers (5,190,838- 5,562,776). However, Graphtyper had the longest compute time (154.1 hours), as the time of constructing and indexing a graph is relatively long for only a single sample.

**Table 3:**
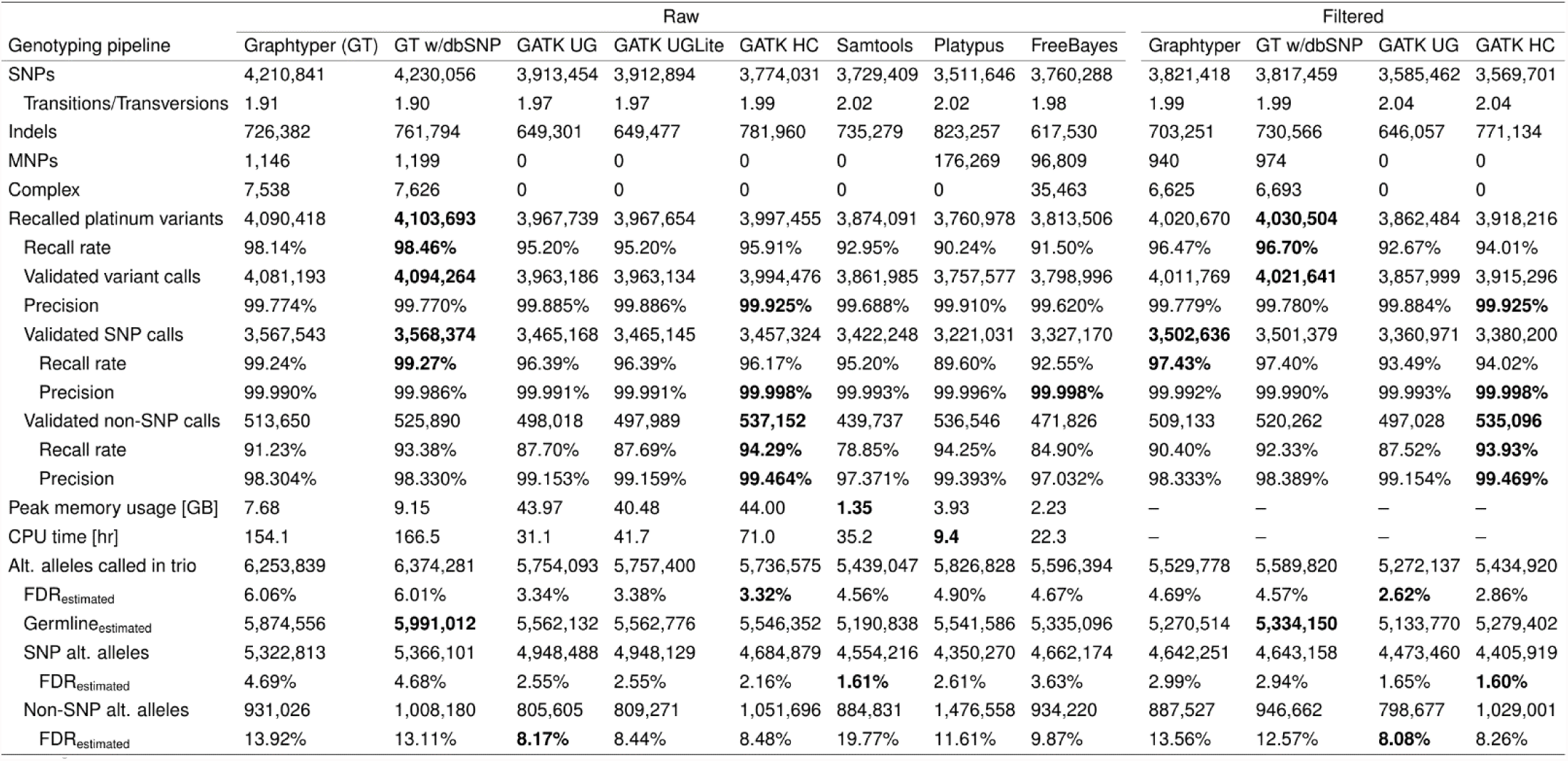
Comparison of whole-genome sequence variant calls of NA12878. Graphtyper was run with and without the knowledge of common dbSNP variation.

We also filtered the Graphtyper call sets (Supplementary Note 4) and compared it with GATK’s call sets filtered according their ‘best practices’ guidelines. After filtering, Graphtyper’s recall rate was reduced to 96.47% and its estimated FDR reduced from 6.06% to 4.69% (Table 3).

### 28,075 Icelandic whole-genome samples

We used Graphtyper to genotype the autosomes and chromosome X of 28,075 whole-genome sequenced Icelandic samples. The samples have a mean sequencing depth of 35.3x (s.d. 7.9x; range 2-200x) stored in a total of 2.12 PB of BAM files. The overall compute time for genotyping was 97,917 CPU days or 83.7 CPU hours per sample on average. Graphtyper genotyped 89.4 million sequence variants: 1.1 million complex variants, 6.4 million indels, and 81.9 million SNPs with a Ti/Tv ratio of 1.04.

The compute time of genotyping chromosome 21 in 28,075 Icelandic samples was 27,853 CPU hours or 0.99 CPU hours per sample on average. Compared to Graphtyper’s chromosome 21 genotyping of 691 samples, the sample size 40-folded, the number of sequence variants increased by 220%, but the compute time per sample only increased by 17.6%.

### HIA typing

The IPD-IMGT/HLA database^8^ contains known HLA allele sequences identified with a field (usually two digits) hierarchical colon separated identifier. The first field denotes the HLA allele family, the second field denotes the subtype within the family, the third field denotes groups with synonymous substitutions within the subtype, and the fourth field denotes allele differences in non-coding regions.

Based on known HLA allele sequences, we created graphs for six important HLA genes: HLA-A, HLA-B, HLA-C, HLA-DRB1, HLA-DQA1, and HLA-DQB1 (Methods). Using these graphs, we were able to HLA type the same dataset of 28,075 Icelanders in a single genotyping-only iteration. Our results show high diversity of HLA allele families in the Icelandic population (Supplementary Table 3).

The total compute time of the HLA genotyping of the six genes was 2,609 hours, or 5.6 minutes per sample. The compute time of Graphtyper for the HLA region was orders of magnitudes lower than other genotypers^32,13^ (Supplementary Note 6). Previously, deCODE genetics laboratory performed HLA typing of the six genes with a PCR based method at 2- digit (*n* = 647) and 4-digit (*n* = 368) resolutions. These previous typings are in good concordance (95.1-100% 2-digit; 91.6-100% 4-digit) with Graphtyper’s HLA genotype calls (Table 4). Upon manual inspection, we concluded that a large fraction of the discrepancy between the two methods are most likely explained by sample mix-up (Supplementary Note 6).

**Table 4:** Comparison of Graphtyper’s HLA typings to PCR verified HLA types.

| HLA gene | *n* | 4 digit resolution |  |  |  | 2 digit resolution |  |  |  |
| --- | --- | --- | --- | --- | --- | --- | --- | --- | --- |
|  |  | Correct | 1 error | 2 errors | Accuracy | Correct | 1 error | 2 errors | Accuracy |
| *HLA-A* | 54 | 52 | 2 | 0 | 98.15% | 52 | 2 | 0 | 98.15% |
| *HLA-B* | 332 | — | — | — | — | 314 | 15 | 3 | 96.84% |
| *HLA-C* | 315 | — | — | — | — | 290 | 19 | 6 | 95.08% |
| *HLA-DQA1* | 42 | 42 | 0 | 0 | 100% | 42 | 0 | 0 | 100% |
| *HLA-DQB1* | 82 | 80 | 2 | 0 | 98.78% | 81 | 1 | 0 | 99.39% |
| *HLA-DRB1* | 190 | 163 | 22 | 5 | 91.58% | 189 | 1 | 0 | 99.74% |

## Discussion

Previous genotypers use read alignments to linear reference genomes, which limits their performance in polymorphic regions. To better characterize sequence diversity we implemented a novel variation-aware data structure and developed efficient algorithms in a software called Graphtyper. Graphtyper locally realigns sequence reads from a genomic region to a pangenome graph, and concomitantly genotypes sequence variants in all individuals. We show that combining these two steps is not only practical, but improves sensitivity and is more scalable than other genotyping methods. Our results show that Graphtyper has the highest Mendelian accuracy at previously reported variant sites among the genotypers in our comparison.

Graphtyper can use known variants as input, further improving sensitivity. When using dbSNP as part of the input, Graphtyper fails to recall only 0.73% of SNP variants in the Platinum genome dataset, a rate 5 times lower than the 3.61% missed by the best competitor. Additionally, the graph representation allows us to construct graphs with known sequence variation in the HLA region and accurately genotype known alleles of six HLA genes. Our HLA types are in good concordance to previously PCR verified HLA types. Graphtyper’s ability to determine genotype calls for more sequence variants, including those that have complex representation, such as the HLA region may help geneticists in characterizing genomes and their impact. Despite these successes, additional work is required, for example, currently Graphtyper cannot call structural variants.

The computational requirements of many genotypers are so large that it is infeasible to effectively apply them to population-sized data sets. For large datasets, the computational requirements of Graphtyper are significantly lower than previous methods, requiring full utilization of a 10,000 core computer cluster for 10 days, compared to an estimated minimum of 25 days for GATK UG.

It is important to note that our current pipeline still relies on the linear reference sequence and BWA for global read alignments in order to assign reads to a region. To completely remove bias towards the reference genome and fully utilize the promise of pangenome analysis requires developing robust methods for graph alignment, some of which are on the horizon ^24,25,27^; one such notable project is vg (https://github.com/vgteam/vg). Our results further show the importance of replacing the linear reference with richer data structures to improve our understanding of how sequence diversity impacts diseases and other phenotypes.

## Methods

### Icelandic DNA data

The Icelandic samples were whole-genome sequenced at deCODE genetics^2^ using Illumina HiSeq and HiSeqX sequencing machines^33^ and aligned to the GRCh38 human reference genome using the BWA MEM algorithm^9^. All sequenced individuals were also SNP chip typed using Illumina Human Hap or Omni chip arrays. DNA was isolated from both blood and buccal samples.

All participating subjects signed informed consent. Personal identities of the participants and biological samples were encrypted by a third party system approved and monitored by the Data Protection Authority. Approvals for these studies were provided by the National Bioethics Committee and the Data Protection Authority in Iceland.

### Sequence read alignment

In Graphtyper, sequence variation of small genomic regions (we used 50 kbp regions this study) are represented with a pangenome graph structure. Sequence reads are realigned to the graph of a region if BWA reported them to be in the same region. First, Graphtyper extracts a set of *k*-mers from the sequence read, which overlap by one DNA base in the read (Figure 3d), and determines if they are present in the graph using an index structure (Figure 3e). Seeds are generated from matches in the index look-up. If the alignments of two adjacent *k*-mers overlap by exactly one base, Graphtyper joins their matches into larger seeds (Figure 3g). The longest seeds are then extended (Figure 3h) by finding a path in the graph with the fewest mismatches using a breadth first search algorithm. If no seeds are extended with 12 mismatches or fewer, Graphtyper again extracts a set of *k*-mers from the read which overlap by one base in a read, but now also *k*-mers with one mismatch are included (Figure 3f). The process is applied both to a read and its reverse complement. If both orientations of a read align to the graph, Graphtyper selects the longer alignment or, if they are equally long, the alignment with fewer mismatches.

### Novel variant discovery

Graphtyper post-processes graph alignments to discover novel small sequence variants. Novel sequence variants are classified as SNPs, indels (up to approx. 50 bp), and complex variation (e.g. multiple nucleotide polymorphisms and microsatellites). For each read uniquely aligned to the graph, Graphtyper starts by determining the position in the reference genome of its first and last aligned position in the graph and extracts the reference sequence between these two positions. Then on each side of the reference sequence, the read is extended by an additional 50 bases plus the number of soft clipped bases on the given side. The read is then locally aligned to the extracted reference sequence using a banded semi-global version of Gotoh’s algorithm (Supplementary Figure 4a). Differences in the local alignments are treated as observations of variants (Supplementary Figure 4b).

Once all reads have been processed, Graphtyper outputs sequence variants where there exists a sample that has at least 5 observations of an alternative allele and its frequency is at least 20% (default values).

### Genotyping

Graphtyper genotype calls sequence variants in the graph by treating the graph alignments as independent observations of each sample’s underlying genotype. It genotypes sequence variants in the graph by considering nearby variants together. Given graph-aligned sequence reads of a population, the likelihood that the reads were sampled from a pair of haplotypes is estimated for each sample and the haplotypes with the highest likelihood are determined. To greatly reduce the number of haplotypes considered, all sequence variants located 5 bp or less from each other are grouped (Supplementary Figure 5a) and each variant group is genotyped independently. Let *H_i_* = { *h*_*i*,1_,*h*_*i*,2_} be a multiset of the unknown haplotypes of sample *i* in a variant group, v, and let 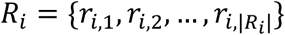 be the sample’s multiset of sequence reads aligned by Graphtyper to the variant group v.

For each pair of possible haplotypes, a relative likelihood of the observed reads given the haplotypes 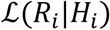 is computed. We assume that the reads from one individual are independent of other individuals’ reads. Graphtyper computes the relative likelihood iteratively as:

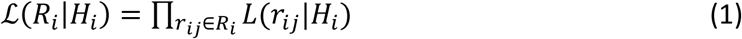

where the relative likelihood of observing a read *r_i,j_* given the pair of underlying haplotypes is set as:

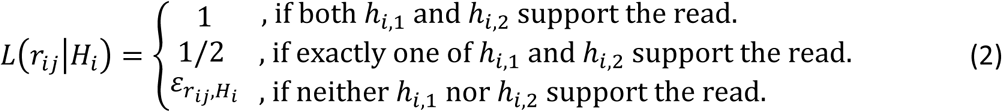

where 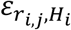 is the relative likelihood of observing an error, given the underlying haplotypes *H_i_* and the read *r_i,j_*. These relative likelihoods are chosen from the set 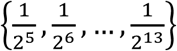 based on how similar the read is to the haplotypes *H_i_*, the base pair quality, mapping quality of the read, and if the read is soft clipped (Supplementary Note 3). Restricting relative likelihoods to this set allows storing only the integer exponents, minimizing storage requirements and avoiding floating point precision problems.

As sequence variants are genotyped in groups, Graphtyper can identify the haplotypes in the population within each group (Supplementary Figure 5b) and remove unobserved haplotypes from the graph (Supplementary Figure 5c). This greatly reduces the number of haplotypes in complex regions.

### Sequence variant quality assessment

For each sequence variant we estimated the Mendelian error rate as the fraction of incorrectly inferred offspring in trios with two homozygous parents (Supplementary Figure 6a). We defined Mendelian inaccuracy as the estimated Mendelian error rate plus the fraction of trios with a missing genotype call, which are genotypes reported as “.” or “./.” in the VCF output.

If either parent is heterozygous we cannot deterministically infer the genotype of the offspring (Supplementary Figure 6b). For those trios we instead calculated the transmission rate of each alternative allele from parent to offspring. The expected transmission rate of germline alternative alleles is 50%. Falsely discovered variation due to sequencing errors and somatic mutations are assumed to transmit at a lower rate. We used the difference of alternative allele transmission rates above and below 50% to estimate the false discovery rate (FDR) using:

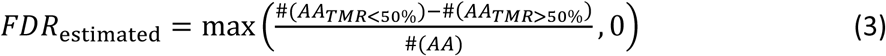

Here, #(*AA*) is the number of called alternative alleles, and #(*AA_TMR_*_>50%_) and #(*AA_TMR_*_<50%_) are the number of alternative alleles with a transmission rate above and below 50%, respectively. We estimated the number of germline alternative alleles using:

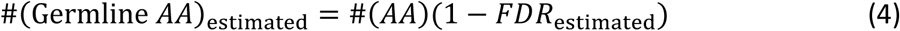

### HLA typing pre-processing

We retrieved HLA allele sequences from the IPD-IMGT/HLA database (version 3.23.0, see URLs). We extracted the differences to a VCF file that we used to create the pangenome graphs for HLA typing. A more detailed description of our HLA typing method as well as comparisons to other methods have been published in our previous work^34^ and are described in Supplementary Note 7.

### Author contributions

HPE implemented the Graphtyper software. HPE, PM and BVH designed the Graphtyper algorithm. HPE, DFG, PM, BVH and KS designed the experiments. HPE, EH, GM and FZ ran all evaluated genotypers. HPE and HJ analyzed the call sets. Aslaug Jonasdottir, Adalbjorg Jonasdottir and IJ were responsible for PCR validation. HJ and SK contributed software for the project. HPE wrote the initial version of the manuscript, HJ, SK, BK, PM, BVH and KS contributed to subsequent versions. All authors reviewed and approved the final version of the manuscript.

### URLs

IPD-IMGT/HLA (http://www.ebi.ac.uk/ipd/imgt/hla/, Github page: https://github.com/ANHIG/IMGTHLA)

### Code availability

Graphtyper is available at https://github.com/DecodeGenetics/graphtyper (GNU GPLv3 license).

